# Vineyard ecosystems are structured and distinguished by fungal communities impacting the flavour and quality of wine

**DOI:** 10.1101/2019.12.27.881656

**Authors:** Di Liu, Qinglin Chen, Pangzhen Zhang, Deli Chen, Kate S. Howell

## Abstract

The flavours of foods and beverages are formed by the agricultural environment where the plants are grown. In the case of wine, the location and environmental features of the vineyard site imprint the wine with distinctive aromas and flavours. Microbial growth and metabolism play an integral role in wine production from the vineyard to the winery, by influencing grapevine health, wine fermentation, and the flavour, aroma and quality of finished wines. The mechanism by which microbial distribution patterns drive wine metabolites is unclear and while flavour has been correlated with bacterial composition for red wines, bacterial activity provides a minor biochemical conversion in wine fermentation. Here, we collected samples across six distinct winegrowing areas in southern Australia to investigate regional distribution patterns of both fungi and bacteria and how this corresponds with wine aroma compounds. Results show that soil and must microbiota distinguish winegrowing regions and are related to wine chemical profiles. We found a strong relationship between microbial and wine metabolic profiles, and this relationship was maintained despite differing abiotic drivers (soil properties and weather/ climatic measures). Notably, fungal communities played the principal role in shaping wine aroma profiles and regional distinctiveness. We found that the soil microbiome is a potential source of grape- and must-associated fungi, and therefore the weather and soil conditions could influence the wine characteristics via shaping the soil fungal community compositions. Our study describes a comprehensive scenario of wine microbial biogeography in which microbial diversity responds to surrounding environments and ultimately sculpts wine aromatic characteristics. These findings provide perspectives for thoughtful human practices to optimise food and beverage flavour and composition through understanding of fungal activity and abundance.

## Introduction

Regional distinctiveness of wine traits, collectively known as “*terroir*”, can be measured by chemical composition and sensory attributes (1–3), and this variation has been related to the physiological response of grapevines to local environments, such as soil properties (e.g., soil type, texture, nutrient availability), climate (temperature, precipitation, solar radiation), topography and human-driven agricultural practices (4–6). Wines made from the same grape cultivar but grown in different regions are appreciated for their regional diversity, increasing price premiums and the market demand (5). However, which factors from the vineyard to the winery drive wine quality traits have remained elusive.

Microbial biogeography contributes to the expression of wine regionality. Microorganisms, including yeasts, filamentous fungi and bacteria, originate in the vineyard and are impacted upon by the built environment (winery) and play a decisive role in wine production and quality of the final product (7, 8). The fermentative conversion of grape must (or juice) into wine is a complex and dynamic process, involving numerous transformations by multiple microbial species (9). The majority of fermentations involve *Saccharomyces* yeasts conducting alcoholic fermentation (AF) and lactic acid bacteria (LAB) for malolactic fermentation (MLF), but many other species are present and impact the chemical composition of the resultant wine (10, 11). Recent studies propose the existence of non-random geographical dispersion patterns of microbiota in grapes and wines (12–18). Few studies have explored the associations between microbial communities and wine chemical composition (19, 20). Knight et al. (20) showed empirically that regionally distinct *S. cerevisiae* populations drove different metabolites present in the resultant wines. Bokulich et al. (19) suggested that wine metabolite profiles were more closely associated with bacterial profiles of wines undergone both AF and MLF. However, both numerically and biochemically, yeasts dominate wine fermentation, so it is unclear why bacteria should be dominant.

The vineyard soil has long been attributed to fundamental wine characteristics and flavour. Vineyard soil provides the grapevine with water and nutrients and soil type and properties profoundly affect vine growth and development (5). Soil-borne microbiota associate with grapevines in a beneficial, commensal, or pathogenic way, and determine soil quality, host growth and health. For example, soil microbes can mineralize soil organic matter, trigger plant defence mechanisms, and thus influence flavour and quality of grapes and final wines (21, 22). Alternatively, serving as a reservoir for grapevine-associated microbiota, the soil microbiome can be recruited from the soil to the grapevine and transferred to the grape and must (14, 23). Some of these microbes can influence the fermentation and contribute to final wine characteristics (8, 24). Overall, biogeographic boundaries can constrain the vineyard soil microbiota (23, 25–28), but correlations between geographical structure of soil microbiota and wine attributes are weak.

Limited but increasing evidence reveals that environmental heterogeneity conditions microbial biogeography in wine production on different spatial scales [recently reviewed by Liu et al. (29); (12, 23, 25, 30, 31). Local climatic conditions significantly correlate with microbial compositions in grape musts, for example precipitation and temperature that correlate with the abundance of filamentous fungi (for example *Cladosporium* and *Penicillium* spp.) and ubiquitous bacteria (for example, the *Enterobacteriaceae* family) (12), as well as yeast populations (particularly *Hanseniaspora* and *Metschnikowia* spp.) (30). Dispersal of soil microbiota is driven by soil physicochemical properties such as soil texture, soil pH, carbon (C) and nitrogen (N) pools (23, 25, 26), with some influences from topological characteristics (for example orientation of the vineyard) (25, 32). Soil microbiome/bacteria may colonise grapes by physical contact (moving by rain splashes, dust, winds) (23) or migration through the plant (xylem/phloem) from the rhizosphere to the phyllosphere (33, 34). Insects help the movement and dispersal of microbes in the vineyard and winery ecosystem, for example, honey bees, social wasps and *Drosopholid* flies can vector yeasts among different microhabitats (35–37). The vineyard microbes enter the winery associated with grapes or must, so the effects of environmental conditions are finally reflected on microbial consortia in wine fermentation. How environmental conditions modulate microbial ecology from the vineyard to the winery and shape regional distinctiveness of wine is still largely unknown.

Here, we sampled microbial communities from the vineyard to the winery across six geographically separated wine-producing regions in southern Australia to provide insights into microbial biogeography, growing environments and wine regionality. We evaluated the volatile chemicals of wines made with Pinot Noir grapes to validate that these different regions have differently flavoured wines. Using next-generation sequencing (NGS) to profile bacterial and fungal communities, we demonstrate that the soil and must microbiota exhibit distinctive regional patterns, and this correlated to the wine metabolome. Network analysis supports microbial community composition that correlated with abiotic factors (weather and soil properties) drives wine regionality both directly and indirectly. The important contributions arise from fungal diversity. We investigated a potential route of wine-related fungi from the soil to the grapes by isolating yeasts from the xylem/phloem of grapevines. Using vineyards, grapes and wine as a model food system, we have related the regional identity of an agricultural commodity to biotic components in the growing system to show the importance of conserving regional microbial diversity to produce distinctive foods and beverages.

## Materials and methods

### Sample sites and weather parameters

15 *Vitis vinifera* cv. Pinot Noir vineyards were selected in 2017 from Geelong, Mornington Peninsula (Mornington), Macedon Ranges, Yarra Valley, Grampians and Gippsland in southern Australia, ranging from 5 km to 400 km between vineyards (Supplementary Figure S1). In 2018, the sampling from the five vineyards in Mornington Peninsula (all < 20 km apart) was repeated to elucidate the influence of sampling year (vintage) on microbial patterns and wine profiles. Each site’s GPS coordinates (longitude, latitude, altitude) were utilised to extract weekly weather data from the dataset provided by Australian Water Availability Project (AWAP). Variables were observed by robust topography resolving analysis methods at a resolution of 0.05° × 0.05° (approximately 5 km x 5 km) (38). Weekly measurements for all vineyards were extracted for mean solar radiation (MS), mean high temperature (MHT), mean low temperature (MLT), maximum temperature (MaxT), minimum temperature (MinT), mean temperature (MT), precipitation, mean relative soil moisture, mean evaporation (soil + vegetation), and mean transpiration (MTrans) in growing seasons (October 2016/2017 - April 2017/2018).

### Collection of soil, plant, must and ferment samples

In each vineyard, soil samples were collected from three sites at harvest March-April 2017 (n = 45) and 2018 (n = 15), at a depth of 0 - 15 cm, 30 – 50 cm from the grapevine into the interrow (Supplementary Table S1A). To further investigate fungal ecology in the vineyard, comprehensive vineyard samples (n = 50) were collected from two vineyards 5 km apart in the 2018 vintage (Supplementary Table S1B). These two vineyards were managed by the same winery, and the viticultural management was very similar, for example, grapevines were under vertical shoot positioning trellising systems and the same sprays were applied at the same time of year. Five replicate Pinot Noir vines in each vineyard were selected from the top, middle, and bottom of the dominant slope, covering topological profiles of the vineyard. For each grapevine, five different sample types were collected: soil (0 - 15 cm deep, root zone), roots, xylem/phloem sap, leaves, and grapes were collected at harvest in March 2018. Xylem sap (n = 10) was collected from the shoots using a centrifugation method in aseptic conditions (39) (Supplementary Table S1B). Details of xylem sap collection, nutrient composition analysis, and yeast isolation were provided in supplementary materials. Samples were stored in sterile bags, shipped on ice, and stored at −80 °C until processing.

Longitudinal samples to study microbial communities during fermentation were collected at five time points: must (destemmed, crushed grapes prior to fermentation), at early fermentation (AF, less than 10% of the sugar is fermented), at middle of fermentation (AF-Mid, around 50% of sugar fermented), at the end of fermentation (AF-End, following pressing but prior to barrelling), at the end of malolactic fermentation (MLF-End, in barrels) (Supplementary Table S1A). Grapes and must sampled in this study were fermented in the wineries following similar fermentation protocols of three days at a cool temperature (known as cold-soaking) followed by warming up the must, so fermentation could commence. Fermentations were conducted without addition of commercial yeasts and bacteria. Two fermentations from Grampians and Mornington did not complete and were excluded from analysis, giving wine samples from 13 vineyards in the 2017 vintage. Triplicate sub-samples from tanks or barrels were collected and mixed as composite samples. All samples (n = 90) were shipped on ice and stored at −20 °C until processing.

### Soil analysis

Edaphic factors were analysed to explore the effects of soil properties on wine-related microbiota and aroma profiles. Soil pH was determined in a 2:5 soil/water suspension. Soil carbon (C), nitrogen (N), nitrate and ammonium were analysed by Melbourne Trace Analysis for Chemical, Earth and Environmental Sciences (TrACEES) Soil Node, at the University of Melbourne. Total C and N were determined using classic Dumas method of combustion (40) on a LECO TruMac CN at a furnace temperature of 1350°C (LECO Corporation, Michigan, USA). Nitrate and ammonium were extracted with 2 M KCl and determined on a Segmented Flow Analyzer (SAN++, Skalar, Breda, Netherlands) (40).

### Wine volatile analysis

To represent the wine aroma, volatile compounds of MLF-End samples were determined using headspace solid-phase microextraction gas-chromatographic mass-spectrometric (HS-SPME–GC-MS) method (41, 42) with some modifications. Analyses were performed with Agilent 6850 GC system and a 5973 mass detector (Agilent Technologies, Santa Clara, CA, USA), equipped with a PAL RSI 120 autosampler (CTC Analytics AG, Switzerland). Briefly, 10 mL wine was added to a 20 ml glass vial with 2 g of sodium chloride and 20 μL of internal standard (4-Octanol, 100 mg/L), then equilibrated at 35°C for 15□min. A polydimethylsiloxane/divinylbenzene (PDMS/DVB, Supelco) 65 μm SPME fibre was immersed in the headspace for 10 min at 35°C with agitation. The fibre was desorbed in the GC injector for 4 min at 220 °C. Volatiles were separated on an Agilent J&W DB-Wax Ultra Inert capillary GC column (30 m × 0.25 mm × 0.25 μm), with helium carrier gas at a flow rate of 0.7 mL/min. The column temperature program was as follows: holding 40 °C for 10 min, increasing at 3.0 °C/min to 220 °C and holding at this temperature for 10 min. The temperature of the transfer line of GC and MS was set at 240 °C. The ion source temperature was 230 °C. The electron impact (EI) at 70 eV scanning over a mass acquisition range of 35–350 *m/z*. Raw data were analysed with Agilent ChemStation Software for qualification and quantification (43). Volatile compounds (n = 88) were identified in wine samples according to retention indices reference standards and mass spectra matching with NIST11 library. 13 successive levels of standard solution in model wine solutions (12% v/v ethanol saturated with potassium hydrogen tartrate and adjusted to pH 3.5 using 40% w/v tartaric acid) were analysed by the same protocol as wine samples to establish the calibration curves for quantification. Peak areas of volatile compounds were integrated via target ions model. The concentrations of volatile compounds were calculated with the calibration curves and used for downstream data analysis.

### DNA extraction and sequencing

Genomic DNA was extracted from plant and soil samples using PowerSoil™ DNA Isolation kits (QIAgen, CA, USA). DNA extraction from soil and xylem sap followed the kit’s protocols. Wine fermentation samples were thawed and biomass was recovered by centrifugation at 4,000 × *g* for 15 min, washed three times in ice-cold phosphate buffered saline (PBS) with 1% polyvinylpolypyrrolidone (PVPP) and centrifuged at 10,000 × *g* for 10 min (12). The obtained pellets were used for DNA extraction following the kit protocol. For grapevine samples, roots, leaves, grapes (removed seeds and stems), were ground into powder under the protection of liquid nitrogen with 1% PVPP, and isolated DNA following the kit protocol afterward. DNA extracts were stored at −20 °C until further analysis.

Genomic DNA was submitted to Australian Genome Research Facility (AGRF) for amplification and sequencing. To assess the bacterial and fungal communities, 16S rRNA gene V3-V4 region and partial fungal internal transcribed spacer (ITS) region were amplified using the universal primer pairs 341F/806R (44) and ITS1F/2 (45), respectively. The primary PCR reactions contained 10 ng DNA template, 2× AmpliTaq Gold® 360 Master Mix (Life Technologies, Australia), 5 pmol of each primer. A secondary PCR to index the amplicons was performed with TaKaRa Taq DNA Polymerase (Clontech). Amplification were conducted under the following conditions: for bacteria, 95 °C for 7 min, followed by 30 cycles of 94 °C for 30 s, 50 °C for 60 s, 72 °C for 60 s and a final extension at 72 °C for 7 min; for fungi, 95 °C for 7 min, followed by 35 cycles of 94 °C for 30 s, 55 °C for 45 s, 72 °C for 60 s and a final extension at 72 °C for 7 min. PCR products were purified, quantified and pooled at the same concentration (5 nM). The resulting amplicons were cleaned again using magnetic beads, quantified by fluorometry (Promega Quantifluor) and normalised. The equimolar pool was cleaned a final time using magnetic beads to concentrate the pool and then measured using a High-Sensitivity D1000 Tape on an Agilent 2200 TapeStation. The pool was diluted to 5nM and molarity was confirmed again using a High-Sensitivity D1000 Tape. This was followed by 300 bp paired-end sequencing on an Illumina MiSeq (San Diego, CA, USA).

Raw sequences were processed using QIIME v1.9.2 (46). Low quality regions (Q < 20) were trimmed from the 5′ end of the sequences, and the paired ends were joined using FLASH (47). Primers were trimmed and a further round of quality control was conducted to discard full length duplicate sequences, short sequences (< 100 nt), and sequences with ambiguous bases. Sequences were clustered followed by chimera checking using UCHIME algorithm from USEARCH v7.1.1090 (48). Operational taxonomic units (OTUs) were assigned using UCLUST open-reference OTU-picking workflow with a threshold of 97% pairwise identity (48). Singletons or unique reads in the resultant data set were discarded, in addition, chloroplast and mitochondrion related reads were removed from the OTU dataset for 16S rRNA. Taxonomy was assigned to OTUs in QIIME using the Ribosomal Database Project (RDP) classifier (49) against GreenGenes bacterial 16S rRNA database (v13.8) (50) or the UNITE fungal ITS database (v7.2) (51), respectively. To avoid/reduce biases generated by varying sequencing depth, sequences were rarefied to an even depth per sample (the lowest sequencing depth of each batch, that is for the soil, must and wine, soil and plant samples) prior to downstream analysis. Raw data are publicly available in the National Centre for Biotechnology Information Sequence Read Archive under the bioproject PRJNA594458 (bacterial 16S rRNA sequences) and PRJNA594469 (fungal ITS sequences).

### Data analysis

Microbial alpha-diversity was calculated using the Shannon index in R (v3.5.0) with the “vegan” package (52). One-way analysis of variance (ANOVA) was used to determine whether sample classifications (e.g., region, fermentation stage) contained statistically significant differences in the diversity. Principal coordinate analysis (PCoA) was performed to evaluate the distribution patterns of wine metabolome and wine-related microbiome based on beta-diversity calculated by the Bray–Curtis distance with the “labdsv” package (53). Permutational multivariate analysis of variance using distance matrices with 999 permutations was conducted within each sample classification to determine the statistically significant differences with “adonis” function in “vegan” (52).

Significant differences of wine microbiome between fermentation stages were tested based on taxonomic classification using linear discriminant analysis (LDA) effect size (LEfSe) analysis (54) (https://huttenhower.sph.harvard.edu/galaxy/). The OTU table was filtered to include only OTUs > 0.01% relative abundance to reduce LEfSe complexity. This method applies the factorial Kruskal–Wallis sum-rank test (α = 0.05) to identify taxa with significant differential abundances between categories (using all-against-all comparisons), followed by the logarithmic LDA score (threshold = 2.0) to estimate the effect size of each discriminative feature. Significant taxa were used to generate taxonomic cladograms illustrating differences between sample classes.

The co-occurrence/interaction patterns between wine metabolome and microbiota in the soil and must were explored in network analysis using Gephi (v0.9.2) (55). Only top 100 OTUs (both bacteria and fungi) based on the relative abundance were used to construct the network. A correlation matrix was calculated with all possible pair-wise Spearman’s rank correlations between volatile compounds and selected OTUs. Correlations with a Spearman correlation coefficient *ρ* ≥ 0.8 and a *p* < 0.01 were considered statistically robust and displayed in the networks (56).

A Random forest supervised-classification model (57) was conducted to identify the main predictors of wine regionality among the following variables: must and soil microbial diversity (Shannon index), soil properties, and weather. The importance of each predictor is determined by evaluating the decrease in prediction accuracy (that is, increase in the mean square error (MSE) between observations and out-of-bag predictions) when the data is randomly permuted for the predictor. This analysis was conducted with 5,000 trees using the “randomForest” package in R (58). The significance of the model and the cross-validated *R*^2^ were assessed using the “A3” package (ntree = 5000) (59). The structural equation model (SEM) (60) was used to evaluate the direct and indirect relationships between must and soil microbial diversity, soil properties, climate and wine regionality. SEM is an *a priori* approach partitioning the influence of multiple drivers in a system to help characterize and comprehend complex networks of ecological interactions (61). An *a priori* model was established based on the known effects and relationships among these drivers of regional distribution patterns of wine aroma to manipulate the data before modelling. Weather and soil properties were used as composite variables (both Random Forest and SEM) to collapse the effects of multiple conceptually-related variables into a single-composite effect, thus aiding interpretation of model results (60). A path coefficient describes the strength and sign of the relationship between two variables (60). The good fit of the model was validated by the χ2-test (*p* > 0.05), using the goodness of fit index (GFI > 0.90) and the root MSE of approximation (RMSEA < 0.05) (62). The standardised total effects of each factor on the wine regionality pattern were calculated by summing all direct and indirect pathways between two variables (60). All the SEM analyses were conducted using AMOS v25.0 (AMOS IBM, NY, USA).

SourceTracker was used to track potential sources of wine-related fungi within the vineyards (63). SourceTracker is a Bayesian approach that treats each give community (sink) as a mixture of communities deposited from a set of source environments and estimates the proportion of taxa in the sink community that come from possible source environments. When a sink contains a mixture of taxa that do not match any of the source environments, that portion of the community is assigned to an “unknown” source (63). In this model, we examined musts (n = 2) and vineyard sources (n = 50) including grapes, leaves, xylem sap, roots, and soils. The OTU tables were used as data input for modelling using the “SourceTracker” R package (https://github.com/danknights/sourcetracker).

## Results

### Chemical composition/aroma profiles separate wines based on geographic location

Using GC-MS, we analysed the volatile compounds of Pinot Noir wine samples (MLF-End) to represent wine metabolite profiles coming from different growing regions and compared directly to the microbial communities inhabiting the musts from which these wines were fermented. In all, 88 volatile compounds were identified in these wines, containing 48 regionally differential compounds (see Table S2 in the supplementary material). Here we used α- and β-diversity measures to further elucidate wine complexity and regionality, respectively. In 2017, α-diversity varied with regional origins (ANOVA, F = 36.021, *p* < 0.001), with higher Shannon indices observed in the wines from regions of Mornington, Yarra Valley and Gippsland (H = 2.17 ± 0.05) than others (H = 1.94 ± 0.03) (Figure 1A). Overall, wine aroma profiles displayed significant regional differentiation across both vintages based on Bray–Curtis dissimilarity (ADONIS, R^2^ = 0.566, *p* < 0.001) and the clustering patterns became more distinct and the coefficient of determination (R^2^) improved when comparing regional differences in 2017 vintage (ADONIS, R^2^ = 0.703, *p* < 0.001) (Supplementary Table S3). PCoA showed that 74.5% of the variance was explained by the first two principal coordinates in 2017, and on PCo1 there were some wines within regions grouped together (Figure 1B).

**Figure 1.**
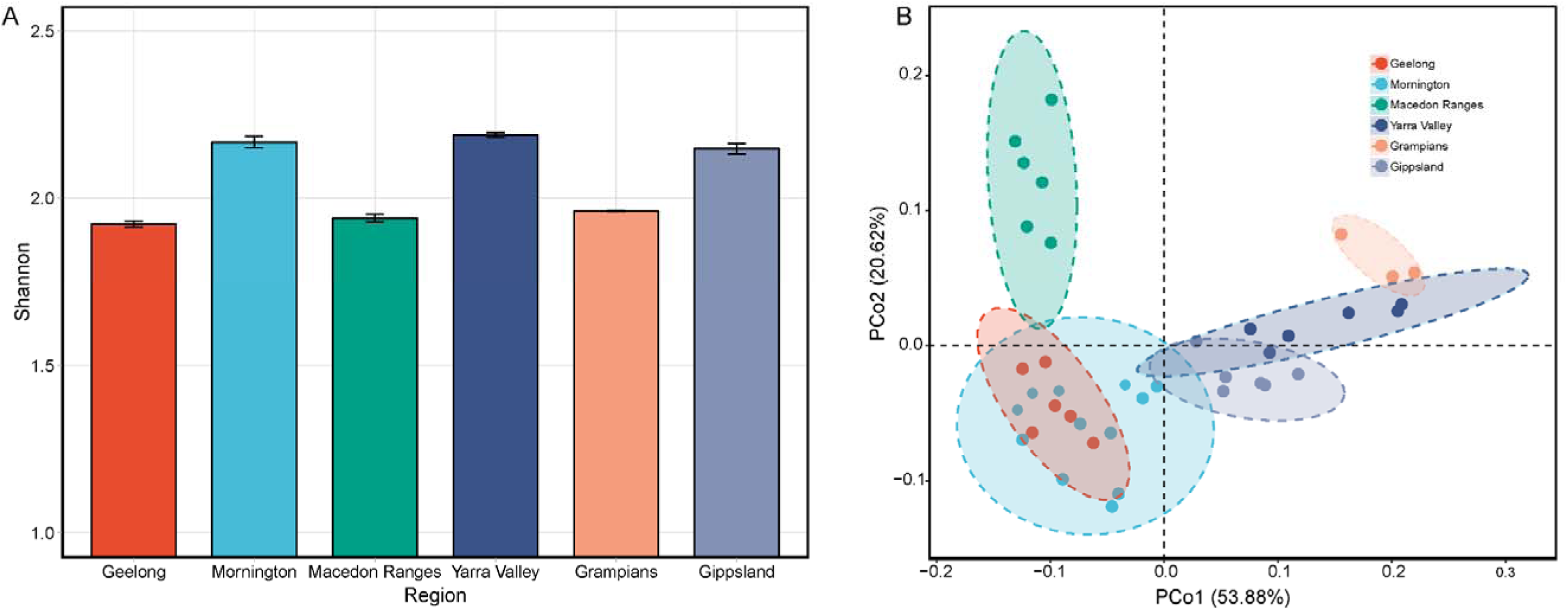
Wine metabolome shows regional variation across six wine growing regions in 2017. Shown are *α*-diversity (Shannon index) (A) and PCoA based on Bray-Curtis dissimilarity obtained from comparing volatile profiles (B).

### Microbial ecology from the vineyard to the winery

To elucidate how microbial ecology drives regional traits of wine from the vineyard to the winery, 150 samples covering soils, musts and fermentations were collected to analyse wine-related microbiota. A total of 11,508,480 16S rRNA and 12,403,610 ITS high-quality sequences were generated from all the samples, which were clustered into 13,689 bacterial and 8,373 fungal OTUs with a threshold of 97% pairwise identity, respectively.

The dominant bacterial taxa across all soil samples were *Actinobacteria*, *Proteobacteria*, *Acidobacteria*, *Chloroflexi*, *Verrucomicrobia*, *Bacteroidetes*, *Gemmatimonadetes*, *Firmicutes*, *Planctomycetes* and *Nitrospirae* (Supplementary Figure S2A). Compared with bacteria, soil fungal communities were less diverse (Supplementary Table S4). *Ascomycota* was the most abundant and diverse phylum of fungi, accounting for 72% of reads, followed by *Basidiomycota*, *Mortierellomycota*, *Chytridiomycota* and *Olpidiomycota* (Supplementary Figure S2B). The microbiome richness (α-diversity, Shannon index) varied significantly with regions for both bacteria and fungi (ANOVA, F_bacteria_ = 4.645, *p* < 0.01; F_fungi_ = 4.913, *p* < 0.01). Soil microbial communities varied widely across different grape growing regions, exerting significant impacts on both bacterial and fungal taxonomic dissimilarity based on Bray-Curtis distances matrices at OTU level (ADONIS, R^2^_bacteria_ = 0.318, *p* < 0.001; R^2^_fungi_ = 0.254, *p* < 0.001), with clearer differences within a single vintage (ADONIS, R^2^_bacteria 2017_ = 0.392, *p* < 0.001; R^2^ = 0.419, *p*< 0.001) (Supplementary Table S3). In 2017, soil fungal communities could discriminate growing regions (except Yarra Valley and Gippsland) (Figure 2B), whereas regional separation of bacteria was weaker with overlap between regions (Figure 2A).

**Figure 2.**
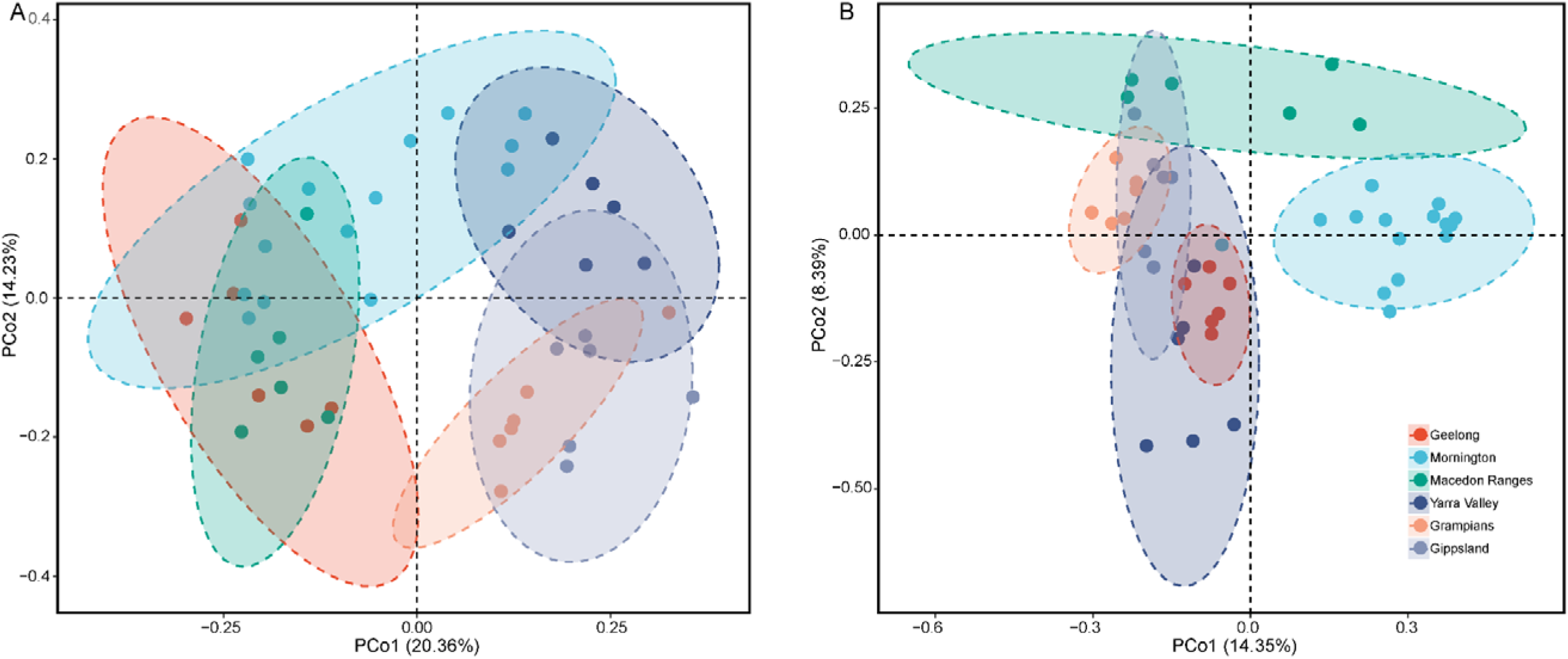
Vineyard soil microbial communities demonstrate regional patterns. Bray-Curtis distance PCoA of bacterial communities (A) and fungal communities (B).

In grape musts, bacterial communities across six wine growing regions in both vintages consisted of ubiquitous bacteria *Enterobacteriales*, *Rhizobiales*, *Burkholderiales*, *Rhodospirillales*, *Actinomycetales*, *Sphingomonadales*, *Pseudomonadales*, *Saprospirales*, and *Xanthomonadales*, which do not participate in wine fermentations or spoilage (7). The LAB *Lactobacillales*, responsible for malolactic fermentation, were present in low abundance (0.4% on average) in the must (Figure 3A). Fungal profiles were dominated by filamentous fungi, mostly of the genera *Aureobasidium*, *Cladosporium*, *Botrytis*, *Epicoccum*, *Penicillium*, *Alternaria*, and *Mycosphaerella*, with notable populations of yeasts, including *Saccharomyces*, *Hanseniaspora*, and *Meyerozyma*, as well as the *Basidiomycota* genus *Rhodotorula* (Figure 3B). Pinot Noir musts exhibited significant regional patterns for fungal communities across vintages 2017 and 2018 based on Bray-Curtis dissimilarity at OTU level (ADONIS, R^2^ = 0.292, *p* < 0.001) but no significant differences for bacterial communities across both vintages (ADONIS, R^2^ = 0.108, *p* = 0.152) (Supplementary Table S3), as well as regarding community richness (ANOVA, F_bacteria_ = 1.567, *p* = 0.374; F_fungi_ = 5.142, *p* < 0.01) (Supplementary Table S4). Within the 2017 vintage, both bacteria and fungi in the must showed distinctive composition based on the region (ADONIS, R^2^_bacteria_ = 0.342, *p* < 0.001; R^2^ = 0.565, *p* < 0.001) (Figure 3A, B) (Supplementary Table S3), with more distinct trend and the improved R^2^ coefficient for fungi. Notably, the relative abundance of *Saccharomyces* yeasts varied significantly from 1.3% (Macedon Range) to 65.6% (Gippsland) between regions (ANOVA, F =26.053, *p* < 0.001) (Figure 3B). As the wine fermentation proceeded, fermentative populations including yeasts and LAB grew and dominated, thus reshaping the community diversity (Supplementary Figure S3A, B) and composition (Supplementary Figure S3C, D). Fungal species diversity collapsed as alcoholic fermentation progressed (ANOVA, F = 6.724, *p* < 0.01) (Supplementary Figure S3B), while the impact of the fermentation on bacterial diversity was insignificant (ANOVA, F = 1.307, *p* = 0.301), with slight decrease at early stages and recovery at fermentation end (Supplementary Figure S3A). Linear discriminant analysis (LDA) effect size (LEfSe) analysis further identified differentially abundant taxa (Kruskal–Wallis sum-rank test, α < 0.05) associated with fermentation stages (Figure 3). For fungal populations, *Dothideomycetes* (including *Aureobasidium* and *Cladosporium*), *Debaryomycetaceae* (notably yeast *M. guilliermondii*), *Penicillium corylophilum* and *Filobasidium oeirense* were significantly abundant in the grape must, with *Leotiomycetes*, *Sarocladium* and *Vishniacozyma victoriae* in early fermentations (AF), *Saccharomycetes* yeasts (notably *S. cerevisiae*) in mid fermentations (AF-Mid), and *Tremellales* at the end of fermentation (AF-End) (Figure 3D). For bacterial communities, *Acidobacteriia* (spoilage), *Chloroflexi*, *Deltaproteobacteria*, *Sphingobacteriia*, *Cytophagia*, *Planococcaceae* and *Rhizobiaceae* were observed with higher abundances in the must, *Proteobacteria* (including *Burkholderiaceae* and *Tremblayales*, spoilage) and *Micrococcus* in the AF, *Burkholderia* spp. in the AF-Mid, *Rhodobacterales* and *Pseudonocardiaceae* in the AF-End, LAB *Leuconostocaceae* (notably *Oenococcus*) in the MLF-End (Figure 3C). Regional differences in microbial profiles were not significant in the finished wines (ADONIS, R^2^_bacteria_ = 0.149, *p* = 0.321; R^2^_fungi_ = 0.109, *p* = 0.205) (Supplementary Table S3).

**Figure 3.**
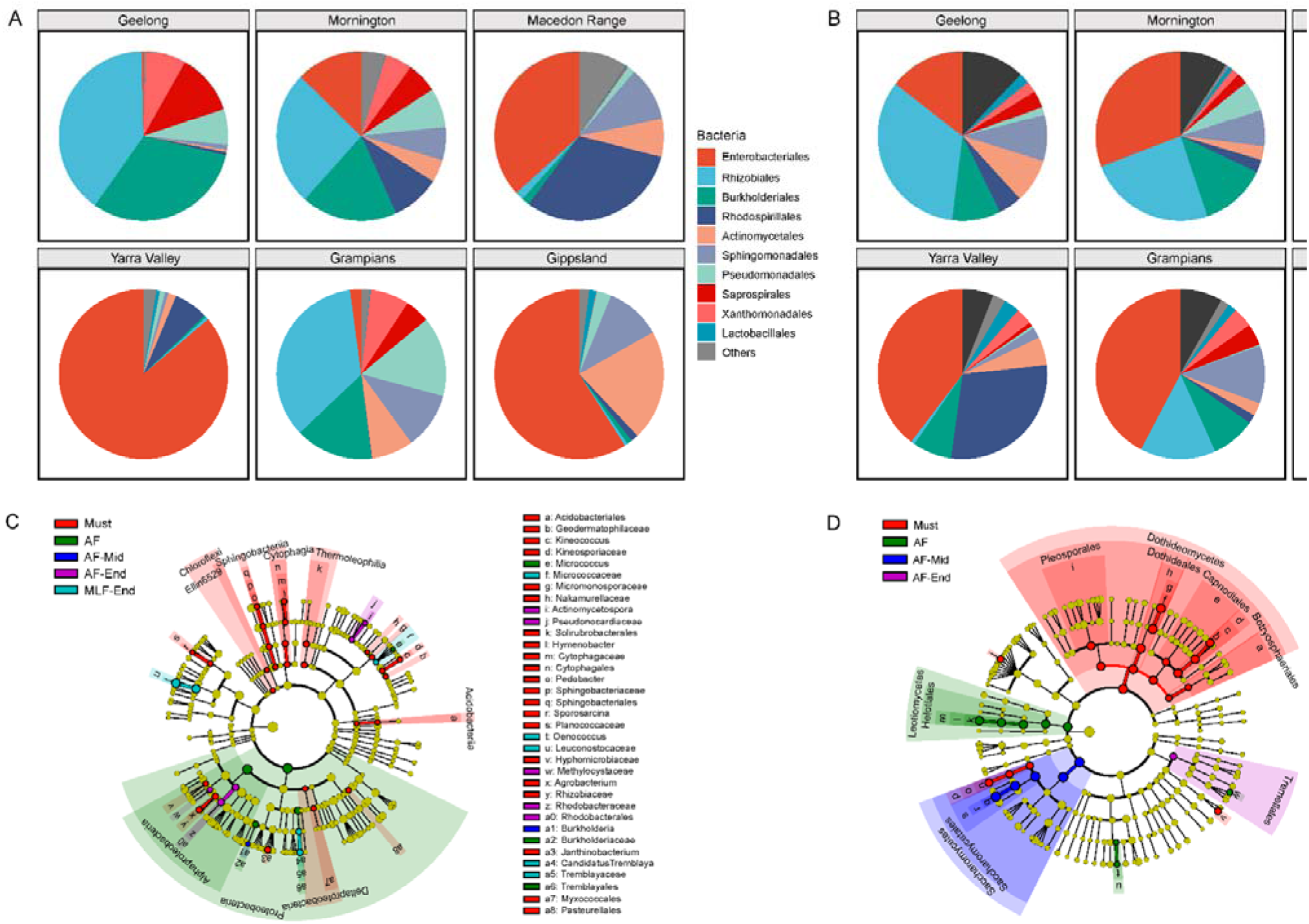
Microbiota exhibit regional differentiation in musts for both bacterial and fungal profiles. Stage of fermentation influences microbial composition of bacteria and fungi. Must bacterial taxa (A) in the order level with greater than 1.0% relative abundance, and *Lactobacillales*. Must fungal taxa (B) in the genus level with greater than 1.0% relative abundance. Linear discriminant analysis (LDA) effect size (LEfSe) taxonomic cladogram comparing all musts and wines categorised by fermentation stage. Significantly discriminant taxon nodes (C, bacteria; D, fungi) are coloured and branch areas are shaded according to the highest-ranked stage for that taxon. For each taxon detected, the corresponding node in the taxonomic cladogram is coloured according to the highest-ranked group for that taxon. If the taxon is not significantly differentially represented between sample groups, the corresponding node is coloured yellow.

To uncover the impact of growing season (vintage) on wine regionality and related microbiota, five vineyards in Mornington were sampled in 2017 and 2018 to compare within and between vintages. Within these five vineyards alone, both microbial communities and wine aroma saw significant influence from vintage effects (Supplementary Table S3). When comparing all samples in the large scale, vintage only weakly impacted microbial and wine aroma profiles, in particular an insignificant influence on fungi (ADONIS, R^2^_fungi_ = 0.049, *p* = 0.066) (Supplementary Table S3). We used 2017 vintage data to further explore microbial biogeography and wine regionality in the following analyses.

### Microbial patterns correlate to wine aroma profiling

Network analysis was used to explore connections between regional microbial and wine metabolic patterns. 57 volatile compounds and 246 OTUs were screened out based on strong correlation coefficients (Spearman correlation coefficient, ρ ≥ 0.8; *p* < 0.01), forming the co-occurrence patterns. These variables showed a sophisticated internal structure, consisting of 303 nodes and 610 edges (average degree = 4.026), with mostly positive correlations (90.66%) (Figure 4A). Both soil (average degree = 3.333; Figure 4B) and must microbiota (average degree = 3.791; Figure 4C) presented high connectivity with wine metabolites (Figure 4B, C). Soil fungal OTUs (69 of 5610 OTUs) more densely connected with volatile compounds than bacteria (42 of 12117 OTUs) (Figure 4B). This indicated that soil microbial communities, especially fungi, might play a role in wine properties. The most densely connected volatile compounds were assigned to groups of monoterpenes (C18, *p*-cymene; C58, α-terpineol), phenylpropanoids (C40, benzaldehyde; C69, phenethyl acetate), and acetoin (C20) (Figure 4A). Correspondingly, high-connectivity nodes with these compounds were OTUs of bacterial taxa *Enterobacteriales*, *Rhizobiales*, *Burkholderiales* and *Xanthomonadales*, and fungal taxa *Penicillium*, *Rhodotorula* and *Neocatenulostroma* from soil and must (Figure 4B, C). The group of esters, which are yeast-driven products (9), were highly associated with fungal OTUs (in particular *Hypocreales*, *Alternaria*, *Cryptococcus and Mortierella*) from the vineyard soil, for example, ethyl hexanoate (C16) and ethyl dihydrocinnamate (C75) (Figure 4B). Fatty acids (including C49, 4-hydroxybutanoic acid; C80, octanoic acid) were mostly negatively connected with fungi (including *Acremonium*, *Penicillium*) from the must (Figure 4C).

**Figure 4.**
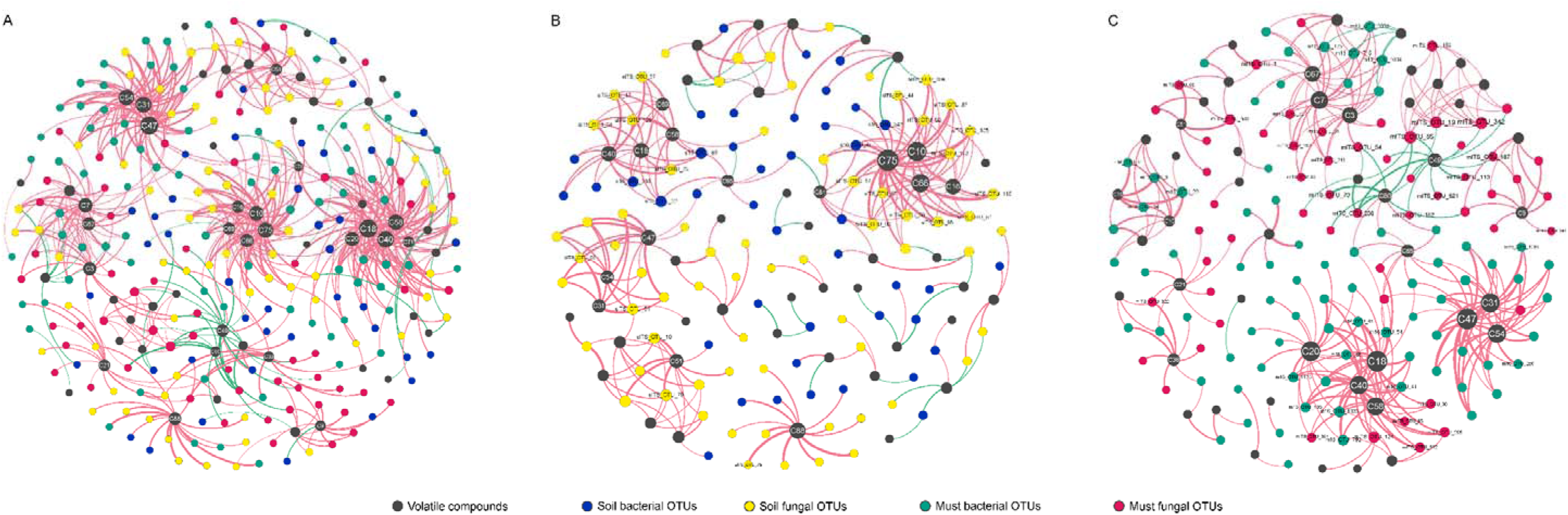
Statistically significant and strong co-occurrence relationships between wine volatile compounds and bacterial and fungal OTUs in vineyard soils and musts across six wine growing regions. Network plots for wine aroma with soil and must microbiota (A), and simplified network plots for wine aroma with soil microbiota (B) and with must microbiota (C), separately. Circle nodes represent assigned volatile compounds, bacterial and fungal OTUs, with different colours. Direct connections between compounds and OTUs indicate strong correlations (Spearman correlation coefficient, *ρ* ≥ 0.8; *p* < 0.01). The colour of edges describes the positive correlation (pink) or the negative correlation (green). The size of nodes is proportioned to the inconnected degree.

### Multiple drivers modify wine regionality in the vineyard

Alongside regional patterns in soil and must microbiota, environmental measures of the wine growing regions displayed significant differences, such as C and N of soil properties, solar radiation and temperature of weather/climatic conditions during the growing season (October 2017 – April 2018) (see Supplementary Table S5 for a complete list). To disentangle the role of microbial ecology on wine regionality, we used Random Forest modelling (57) to identify biotic (soil and must microbial diversity) and abiotic predictors (soil and weather parameters) structuring wine regionality, and structural equation modelling (SEM) (60) to testify whether the relationship between microbial diversity and wine regionality can be maintained when accounting for multiple drivers simultaneously. The Random Forest model (R^2^ = 0.451, *p* < 0.01) demonstrated that fungal diversity was a major predictor of wine regionality. Not surprisingly, must fungal diversity showed higher importance on the model (increase in MSE) than soil (Figure 5). The SEM explained 93.8% of the variance found in the pattern of wine regionality (Figure 6A). Weather drove wine aroma profiles directly (especially MT, MLT, MinT and MS) and indirectly by powerful effects on soil and must microbial diversity, in particular strong effects on soil fungal diversity (Figure 6A). Must fungal diversity had the highest direct positive effect on wine aroma characteristics, with direct influences from soil fungal diversity (Figure 6A). Weather and climate could impact soil nutrient pools primarily through MS, MLT, MinT, and MTrans. Soil properties showed strong effects on soil microbial diversity and must bacterial diversity, but weak effects on must fungal diversity (Figure 6A). Must bacterial diversity had a weak effect on wine aroma profiles, as well as soil bacteria. Overall, must fungal diversity was the most important driver of wine characteristics, followed by soil fungal diversity (Figure 6B), with influences from weather and soil properties both directly and indirectly (Figure 6A).

**Figure 5.**
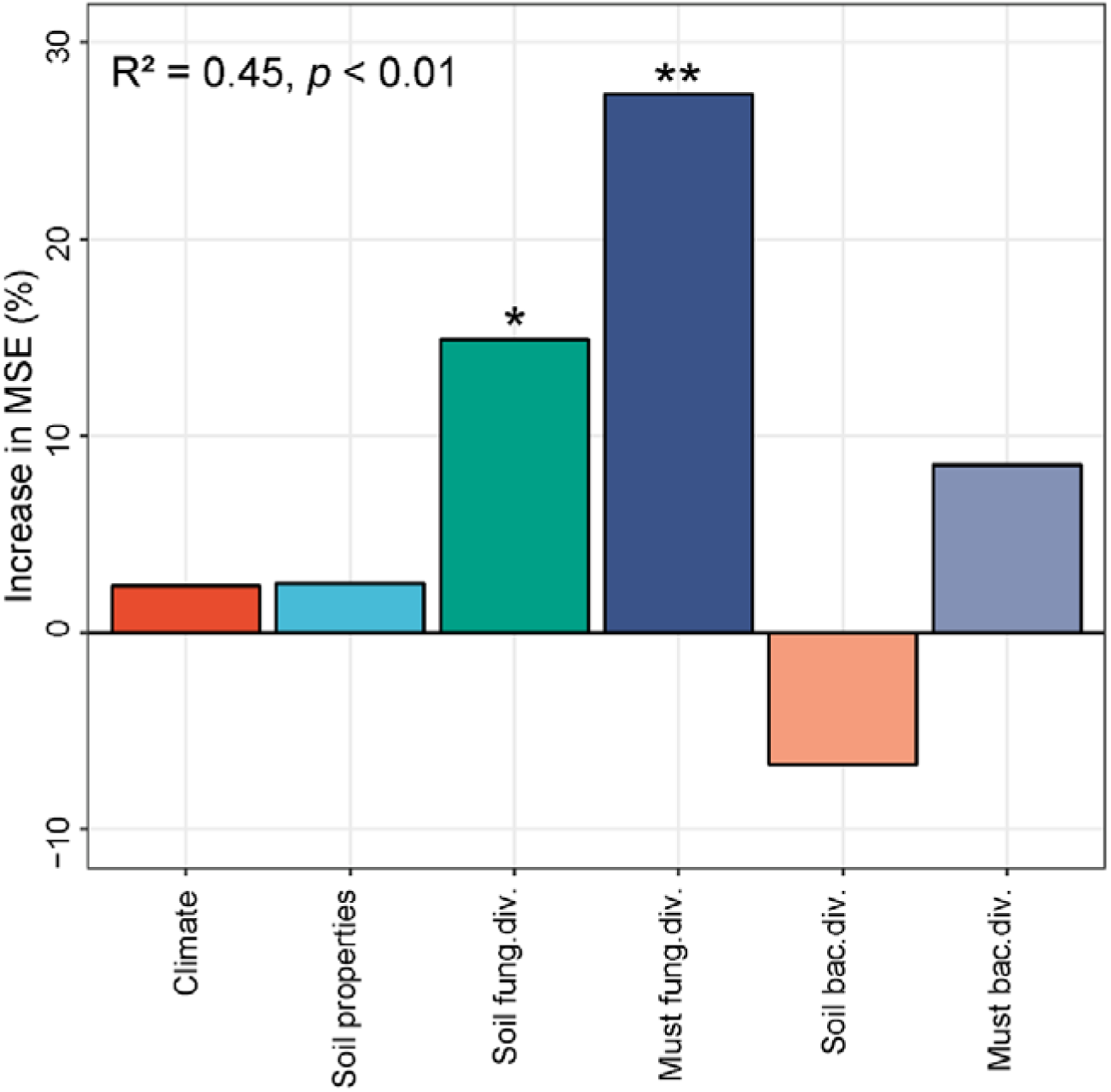
Main predictors of wine regionality. Shown is the Random Forest mean predictor importance by % of increase in mean square error (MSE) of climate, soil properties and microbial diversity (Shannon index) on wine regionality. Soil bac. div., soil bacterial diversity; Soil fung. div., soil fungal diversity; Must bac. div., must bacterial diversity; Must fung. div., must fungal diversity. Significance levels: \**p* < 0.05, \*\**p* < 0.01.

**Figure 6.**
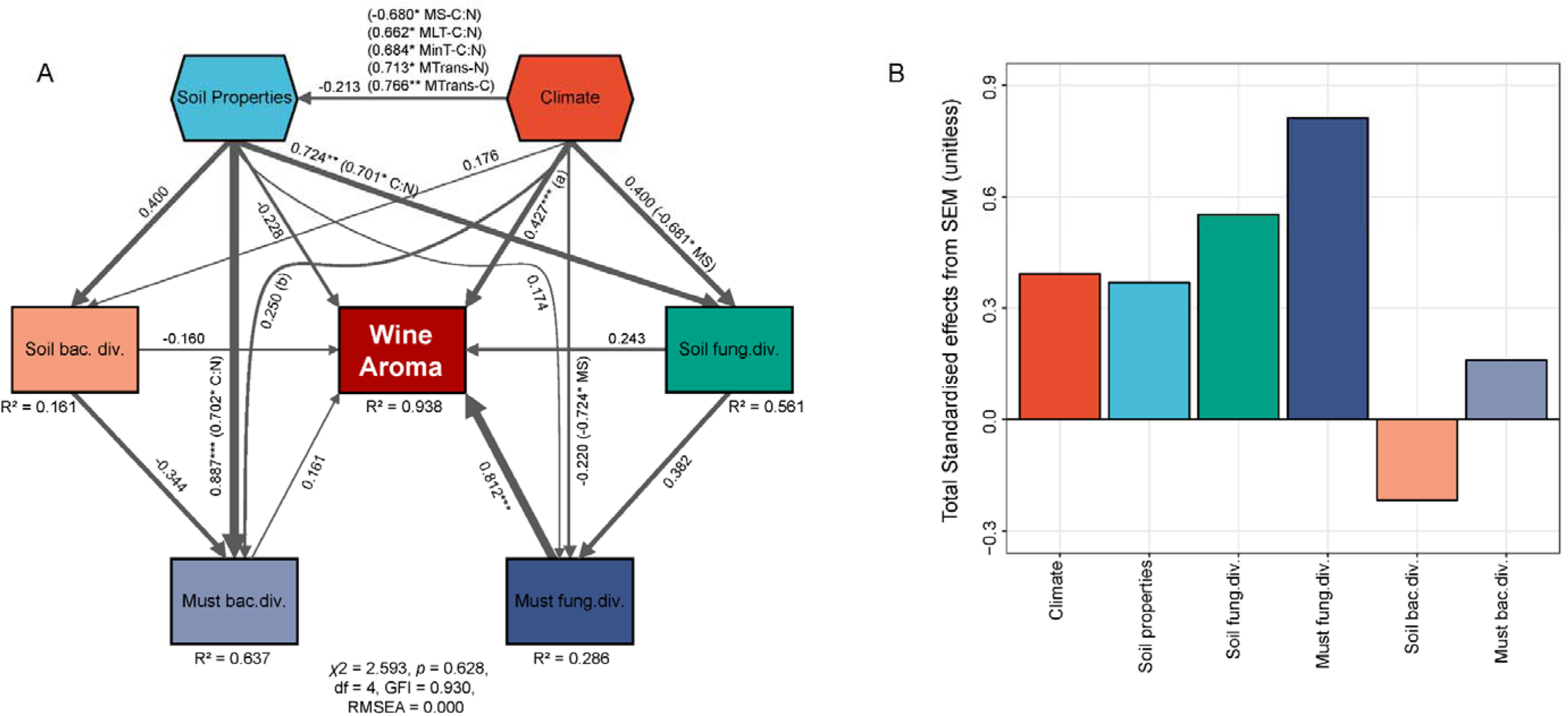
Direct and indirect effects of climate, soil properties and microbial diversity (Shannon index) on wine regionality. Structural equation model (SEM) fitted to the diversity of wine aroma profiles (A) and standardized total effects (direct plus indirect effects) derived from the model (B). Climate and soil properties are composite variables encompassing multiple observed parameters (see materials and methods for the complete list of factors used to generate this model). Numbers adjacent to arrows are path coefficients and indicative of the effect size of the relationship. The width of arrows is proportional to the strength of path coefficients. *R*^2^ denotes the proportion of variance explained. (a) (0.747* MT) (0.666*MLT) (0.686* MinT) (−0.875** MS). (b) (0.753* MinT) (0.772* MLT) (−0.683* MS) (−0.737* MHT) (−0.843** MaxT). C, soil carbon; N, soil nitrogen; C:N, soil carbon nitrogen ratio; MS, mean solar radiation; MT, mean temperature; MLT, mean low temperature; MHT, mean high temperature; MinT, minimum temperature; MaxT, maximum temperature; MTrans, mean transpiration. Significance levels: \**p*< 0.05, \*\**p* < 0.01, \*\*\**p* < 0.001.

### Source tracking wine-related fungi within vineyard

As we showed in the SEM, soil fungal diversity had a direct positive influence on must fungal diversity and thus indirectly contributed to wine aroma profiles (Figure 6). Given that soil is a potential source of fungi associated with wine production (14), here we attempt to uncover the mechanism whereby soil fungi could be transported from soil to the grapes. We sampled fungal communities from grapevines and soil and hypothesised that the xylem/phloem was the internal mechanism to transport microbes. A total of 2,140,820 ITS high-quality sequences were generated from soil and grapevine samples (grape, leaf, xylem sap, root), which were clustered into 4,050 fungal OTUs with 97% pairwise identity. Using SourceTracker (63), fungal communities in the must were matched to multiple sources from below the ground to above the ground. Results showed that grape and xylem sap were primary sources of must fungi, with 32.6% and 41.9% contributions, respectively (Figure 7). Xylem sap showed similar fungal structure with must (Figure S4A). Further source tracking revealed that the root and soil contributed 90.2% of fungal OTUs of xylem sap, the latter contributed 67.9% of the fungi of grapes (Figure 7).

**Figure 7.**
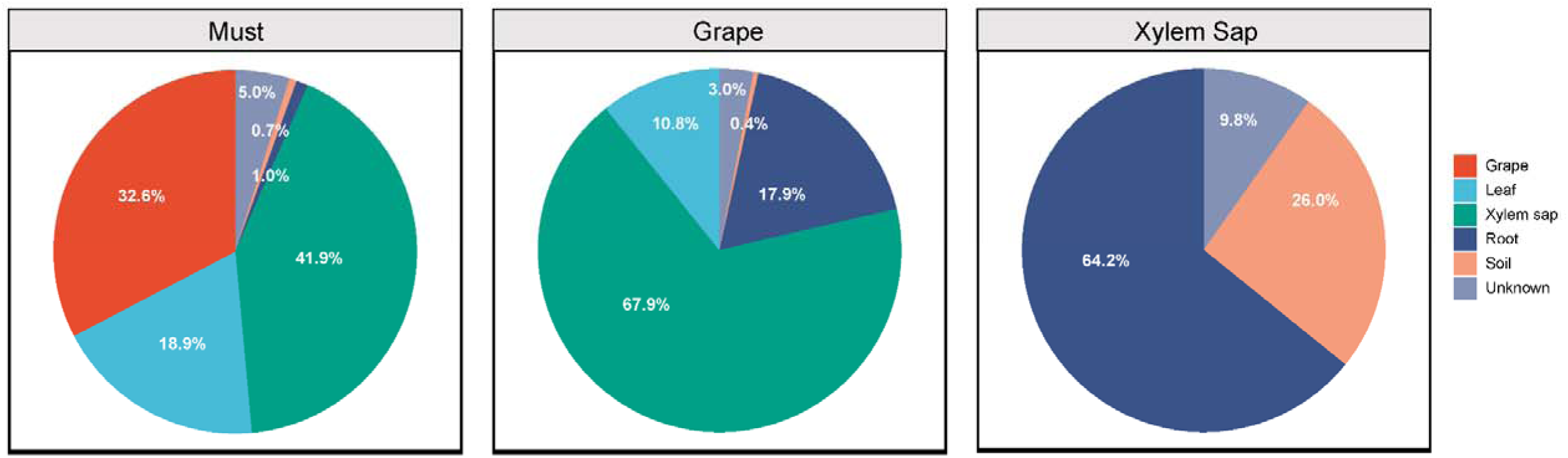
Fungal communities in musts emerge from multiple sources in the vineyard, but primarily from grape and xylem sap. Percent composition estimate for the contributions of possible sources in the

Notably, *S. cerevisiae* were found shared between microhabitats of soil, root, xylem sap, grape and must, with the highest (1.22%) and lowest (0.038%) relative abundance in the root and soil, respectively (Figure S4B). Could xylem vessels be a possible translocation pathway of *S. cerevisiae* from roots to the aboveground? Chemical analysis of nutrient compositions showed that xylem sap contained nine carbohydrates (predominantly glucose, fructose and sucrose), 15 amino acids (mainly arginine, aspartic acid and glutamic acid) and six organic acids (primarily oxalic acid), which could be utilised as carbon sources and support yeast growth (Figure S4E, F, G) (64). However, no *S. cerevisiae* yeasts were isolated, distinct isolates of the *Basidiomycota* yeasts of *Cryptococcus* spp. (primarily *C. saitoi*) and *Rhodotorula slooffiae* were found instead (Figure S4C). The exclusive existence of these species was validated by isolation from xylem/phloem sap coming from grapevines grown in the glasshouse (Figure S4D).

## Discussion

Microbial ecology can influence grapevine health and growth, fermentation, flavour characteristics, and wine quality and style (12, 13, 20). We systematically investigated the microbiome from the soil to wine to show that soil and grape must microbiota exhibit regional patterns and that these patterns correlate with wine metabolites and thus could affect wine quality and style. Here we show that wine regionality is dependent on fungal ecology and is sensitive to local weather, climate and soil properties. A new mechanism to transfer fungi from the soil to grapes and must via xylem sap was investigated.

### A microbial component of wine *terroir*

Regional spatial patterns have been proposed for soil and grape must microbiota (12, 25–28). The most abundant bacterial phyla in the vineyard soils in our study were *Actinobacteria*, *Proteobacteria* and *Acidobacteria*, which are known to be dominant and ubiquitous in vineyard soil (23, 25, 26, 65). For fungi, we recovered 14 phyla, 30 classes, 65 orders, 125 families and 216 genera of fungi, recording a higher diversity than reported in other wine-producing areas in the world (26, 27, 66). *Glomeromycota*, the phylum of arbuscular mycorrhizal fungi reported to positively affect grapevine growth, was reported as abundant in New Zealand vineyards (66). In our study, which used amplicon sequences at the ITS region (rather than the D1/D2 region in (66)), *Glomeromycota* was only recovered with low frequency from Mornington and Macedon Ranges vineyards. Clearly, there are differences based on the barcoding region but geographic location may also affect distribution (67), as Coller et al. (2019) using the same ITS1F/2 primers as in our study, retrieved *Glomeromycota* as a core phylum member from vineyards in Italy (26). In the must, both principal fermentation drivers (*S*. *cerevisiae* and LAB) and innocent species (those that neither ferment nor spoil, such as *Enterobacteriales* and *Aureobasidium*) were present in different abundances among regions (Figure 3A, B). The order *Lactobacillales*, representing LAB, was present 0.4% across regions, compared to 29.7% found in California, US (12) and 14% in Catalonia, Spain (32).

Environmental factors (such as weather and climate) and geographic features structure microbial diversity and biogeography across various habitats in the soil and plant ecosystems (12, 25, 68–70). In this study, we demonstrate that microbial biogeographic communities were distinct in both vineyard soils and grape musts in southern Australia regardless of the growing season/vintage. This aligns with previous studies on wine microbial biogeography and provides further evidence for microbial *terroir* [reviewed by Liu et al.(29); (12, 14, 18, 25)]. Soil bacteria can distinguish wine growing regions, with impacts from soil properties (Figure 6A) and this is supported by previous work in this field (23, 25, 28). An interesting finding is that the must bacterial diversity is strongly influenced by soil properties, in particular C: N. Previous work has shown that structures of must and soil communities are similar and some species *Enterobacteriales* and *Actinomycetales* originate from the soil (8, 15). As C: N can be manipulated by composting and cover crops (71), there is an opportunity to manipulate wine microbiota by a focus on vineyard management (29). Soil bacterial microflora is recognised as important for plant growth processes more broadly (72), but fungal diversity beyond endosymbiotic mycorrhizae has not been systematically investigated for grapevines. Here, we show that the soil fungal communities are distinct for a region. Our modelling suggests that soil properties and weather strongly affect soil fungal diversity, which was in line with large-scale studies in which climatic (especially precipitation) and edaphic factors (especially C: N) are the best predictors of soil fungal richness and community composition (70, 73). Must fungal diversity is also shaped by weather and soil properties indirectly via soil fungi that ultimately affect wine aroma profiles.

It is noteworthy that the drivers of microbial patterns change during wine fermentation. Microbial diversity decreased as alcoholic fermentation proceeds, with a clear loss of microorganisms including filamentous fungi, non-*Saccharomyces* yeasts (for example *M. guilliermondii*), spoilage bacteria *Acidobacteria* and *Proteobacteria*, and other bacteria with unknown fermentative functions (for example *Chloroflexi*) (Figure 3C, D), and thus the biogeographic trend was lost by the stage of MLF-End (Supplementary Table S3). This trend was observed more distinctly in fungal communities compared with bacteria (Figure S3A, B). This is not unexpected as it is clear that fermentation more strongly affects fungal populations compared with bacteria, due to increasing fermentation rate, temperature and ethanol concentration induced by *S. cerevisiae* growth (10, 74). In this case, fermentation conditions, such as the chemical environment and interactions and/or competition within the community (10, 75), reshape the microbial patterns observed. Despite the complex change of microbial ecosystem during fermentation, we show that biogeographic patterns in the must could be reflected in the regional metabolic profiling of wine. Our modelling indicates that indirect influences of weather and soil properties via shaping soil and must microbial diversity are more powerful than the direct influences on wine aroma profiles (Figure 6). In the resulting wines, most volatile compounds were alcohols, esters, acids and aldehydes, some of which were likely microbial products. Some compunds are grape-derived and are modified by microbial metabolism during fermentation (9). Co-occurrence networks demonstrate that the soil and must microbiota correlate closely to the wine metabolome. These interactions also indicate potential modulations of soil and must microbiota on wine chemistry, in particular those taxa that are numerically dominant early in the fermentation. For example, bacteria *Enterobacteriales* positively correlated with some grape-derived volatile compounds (monoterpenes and phenylpropanoids), thus potentially enhancing varietal aroma in wine (9). *Enterobacteriales* have been also previously related to wine fermentation rate by Bokulich et al. (19). Fungal genera abundance (for example *Penicillium*) negatively correlated with the medium-chain fatty acid octanoic acid, which can contribute to mushroom flavour and also alter *Saccharomyces* metabolism (Bisson 1999). These microorganisms correlate with wine metabolites and/or can change chemical environments and fermentation behaviours and finally exert substantial effects on wine metabolite profiles (9, 19). It is particularly important to inform farming practices to structure regional microbial communities that can benefit soil quality, and thus, crop productivity.

### Fungal communities distinguish wine quality and style

In grape musts, bacterial and fungal communities exhibit different responses to site-specific and environmental effects. Bacterial regional patterns were not as distinct as fungi, and significantly impacted by vintage (Supplementary Table S3). Although profound responding to soil properties (for example C:N) and affecting wine fermentation, must bacteria show insignificant influences on wine aroma profiles (Figure 6). In contrast, fungal communities display discriminating distribution patterns at the regional scale and are weakly or insignificantly impacted by vintage in this study, aligning with results presented by Bokulich et al. (2014) (12). Soil fungal communities are less diverse than bacterial communities (Figure S2; ref. 67), but of more importance to the resultant wine regionality (see Figure 4, 5, 6). Must fungi, in particular the fermentative yeasts, conduct alcoholic fermentations and provide aroma compounds to influence wine flavour (9). As indicated by SEM, soil fungal communities are affected by local soil properties and weather and exert impacts on must fungal communities (Figure 6). Co-occurrence relationships between wine metabolome and soil microbiota highlight potential contributions from fungal taxa (Figure 4). One explanation is that grapevines filter soil microbial taxa selecting for grape and must consortia (76, 77). Beyond fungi, plant fitness is linked strongly to the responses of soil microbial communities to environmental conditions (78). More sensitive responses of vineyard soil fungi might improve grapevine fitness to local environments thus benefiting the expression of regional characteristics of grapes and wines.

How could yeasts present in the soil be transported to the grape berry? Soil is a reservoir of grapevine-associated microbiome (Figure 7) that is supported by previous publications (14, 23, 28, 79). As well as transporting water and minerals absorbed by roots to the photosynthetic organs, xylem sap is also a microhabitat for microbes that can bear its nutritional environment (64). Here we investigated xylem sap as a conduit to shape the microbiota in the grape by enrichment of the microbes recruited by roots and transported by xylem sap to the grape berries (23, 33, 72). The isolated yeasts belong to the *Cryptococcus* and *Rhodotorula* genera, indicating that the xylem sap environment is not sterile and can potentially transport yeasts to the phyllosphere. The endophyte *Burkholderia phytofirmans* strain PsJN has been shown to colonise grapevine roots from the rhizosphere and spread to inflorescence tissues through the xylem (33, 80). While we were unable to find fermentative yeasts in the sap of grapevines, other yeasts and/or spores were present and thus may also be transported within the grapevine as well as making their way to the phyllosphere through other mechanisms (water splashes, insect vectors). As previous studies show, fermentative yeasts are persistent in vineyards (81, 82) and can be transported by grapevines (34). We can thus conclude that fungi are the principal drivers of consistent expression of regionality in wine production.

Our study illustrates that microbes are absolute contributors to wine aroma and that this comes from the environment in which they are grown and provides a basis to explain how wine may have a distinctive flavour from a particular region. Fungi play a crucial role in interrelating these biotic and abiotic elements in vineyard ecosystems and could potentially be transported internally. Climate and soil properties profoundly structure microbial patterns from the soil to the grape must, which ultimately sculpt wine metabolic profiling. We do not yet know how grapevines recruit their microbiome to maximise physiological development and encourage and maintain microbial diversity under local conditions.

## Acknowledgements

The authors give their sincere thanks to the vignerons who kindly allowed vineyard access, enabled sampling and provided wine samples. DL acknowledges support from a Ph.D. scholarship and funding from Wine Australia (AGW Ph1602) and a Melbourne Research Scholarship from the University of Melbourne.

## Conflict of Interest

The authors declare that the research was conducted in the absence of any commercial or financial relationships that could be construed as a potential conflict of interest.

## Supplementary Tables and Figures

**Table S1.** Number of soil, must and ferments, and plant samples collected in this study.

(A) Soil, must and ferment samples
| Vintage | Region | Vineyard (V) | Sample Fraction |  |  |  |  |  |
| --- | --- | --- | --- | --- | --- | --- | --- | --- |
|  |  |  | Soil | Must<sup>1</sup> | AF<sup>1</sup> | AF-Mid<sup>1</sup> | AF-End<sup>1</sup> | MLF-End<sup>1,2</sup> |
| 2017 | Geelong | V1 | 3 | 1 | 1 | 1 | 1 | 1 |
|  |  | V2 | 3 | 1 | 1 | 1 | 1 | 1 |
|  | Mornington | V3 | 3 | 1 | 1 | 1 | 1 | 1 |
|  |  | V4 | 3 | 1 | 1 | 1 | 1 | 1 |
|  |  | V5 | 3 | 1 | 1 | 1 | 1 | 1 |
|  |  | V6 | 3 | 1 | 1 | 1 | 1 | 1 |
|  |  | V7 | 3 | N/A | N/A | N/A | N/A | N/A |
|  | Macedon Ranges | V8 | 3 | 1 | 1 | 1 | 1 | 1 |
|  |  | V9 | 3 | 1 | 1 | 1 | 1 | 1 |
|  | Yarra Valley | V10 | 3 | 1 | 1 | 1 | 1 | 1 |
|  |  | V11 | 3 | 1 | 1 | 1 | 1 | 1 |
|  | Grampians | V12 | 3 | 1 | 1 | 1 | 1 | 1 |
|  |  | V13 | 3 | N/A | N/A | N/A | N/A | N/A |
|  | Gippsland | V14 | 3 | 1 | 1 | 1 | 1 | 1 |
|  |  | V15 | 3 | 1 | 1 | 1 | 1 | 1 |
| 2018 | Mornington | V3 | 3 | 1 | 1 | 1 | 1 | 1 |
|  |  | V4 | 3 | 1 | 1 | 1 | 1 | 1 |
|  |  | V5 | 3 | 1 | 1 | 1 | 1 | 1 |
|  |  | V6 | 3 | 1 | 1 | 1 | 1 | 1 |
|  |  | V7 | 3 | 1 | 1 | 1 | 1 | 1 |

<sup>1</sup> For microbial analysis: triplicates from tanks/ barrels were collected and mixed as one composite sample which was analysed by NGS
<sup>2</sup> For wine aroma analysis: triplicates were analysed by GC-MS

N/A: not applicable as spontaneous fermentations did not complete and were excluded from analysis (B) Soil and plant samples
| Vintage | Region | Vineyard (V) | Vine (v) | Sample Fraction |  |  |  |  |
| --- | --- | --- | --- | --- | --- | --- | --- | --- |
|  |  |  |  | Soil | Root | Xylem sap | Leaf | Grape |
| 2018 | Mornington | V5 | v1 | 1 | 1 | 1 | 1 | 1 |
|  |  |  | v2 | 1 | 1 | 1 | 1 | 1 |
|  |  |  | v3 | 1 | 1 | 1 | 1 | 1 |
|  |  |  | v4 | 1 | 1 | 1 | 1 | 1 |
|  |  |  | v5 | 1 | 1 | 1 | 1 | 1 |
|  |  | V6 | v6 | 1 | 1 | 1 | 1 | 1 |
|  |  |  | v7 | 1 | 1 | 1 | 1 | 1 |
|  |  |  | v8 | 1 | 1 | 1 | 1 | 1 |
|  |  |  | v9 | 1 | 1 | 1 | 1 | 1 |
|  |  |  | v10 | 1 | 1 | 1 | 1 | 1 |

**Table S2.**
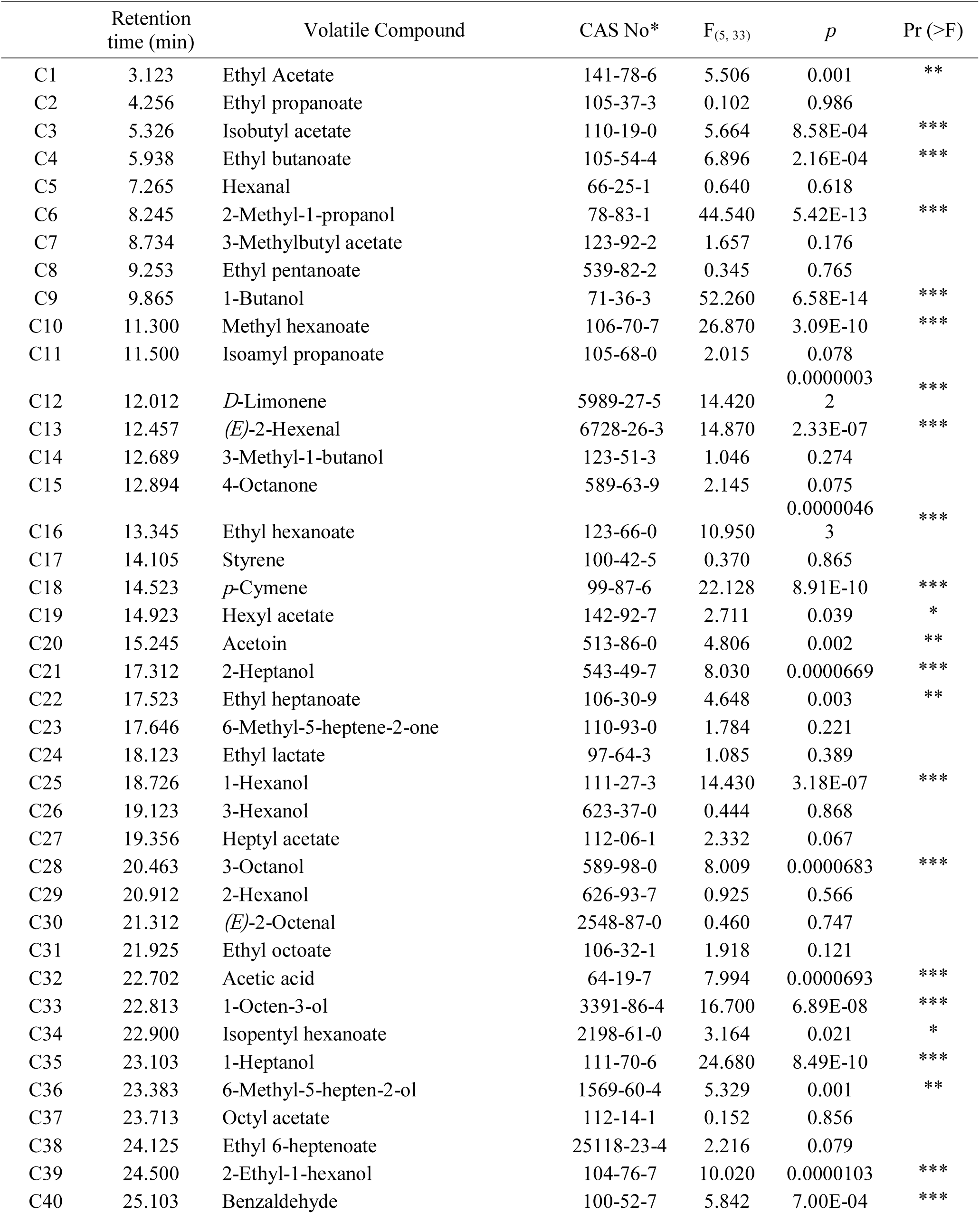
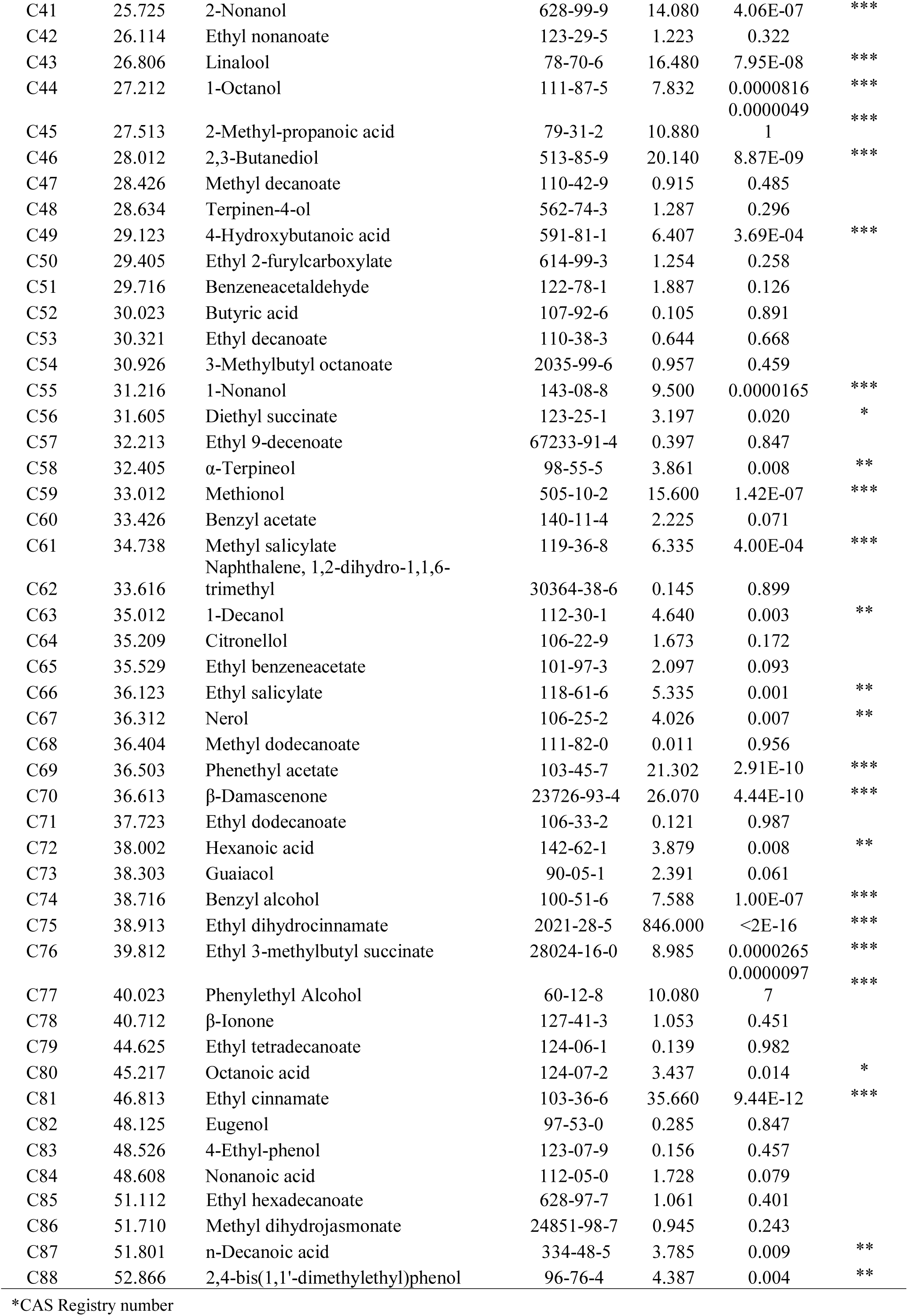
ANOVA results of wine volatile compounds among wine growing regions in 2017.

**Table S3.**
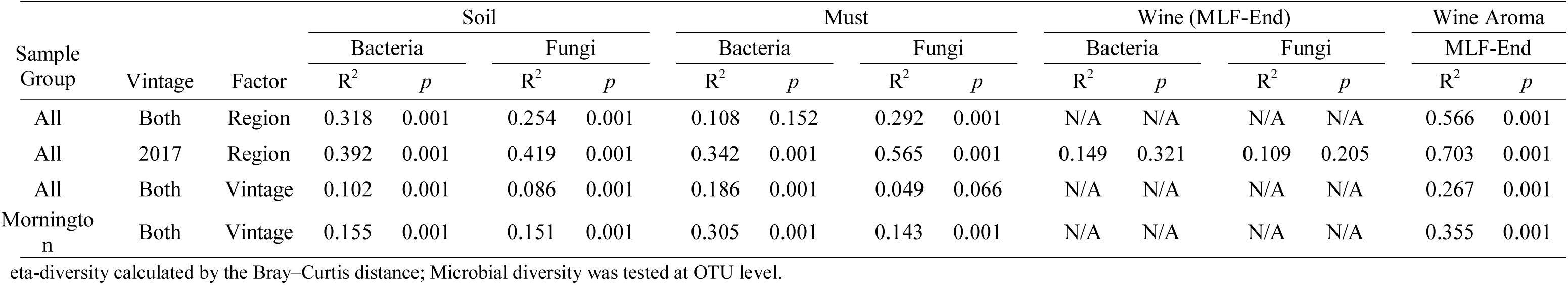
Permutational multivariate analysis of variance using distance matrices of category effects on microbial and wine aroma diversity.

**Table S4.** α-diversity (Shannon index) of soil and must microbial communities from six wine growing regions in 2017 and 2018.

| Vintage | Region | Soil |  | Must |  |
| --- | --- | --- | --- | --- | --- |
|  |  | Bacteria | Fungi | Bacteria | Fungi |
| 2017 | Geelong | $5.486 \pm 0.058$ | $3.869 \pm 0.181$ | $1.838 \pm 0.145$ | $0.938 \pm 0.125$ |
| | Mornington | $6.178 \pm 0.074$ | $4.472 \pm 0.139$ | $2.216 \pm 0.242$ | $1.840 \pm 0.217$ |
| | Macedon Ranges | $4.317 \pm 0.047$ | $2.654 \pm 0.148$ | $2.424 \pm 0.425$ | $1.582 \pm 0.246$ |
| | Yarra Valley | $6.201 \pm 0.062$ | $3.355 \pm 0.238$ | $1.756 \pm 0.314$ | $2.560 \pm 0.123$ |
| | Grampians | $6.238 \pm 0.120$ | $3.615 \pm 0.191$ | $1.966 \pm 0.125$ | $1.297 \pm 0.086$ |
| | Gippsland | $6.416 \pm 0.036$ | $4.080 \pm 0.227$ | $1.855 \pm 0.245$ | $2.200 \pm 0.361$ |
| 2018 | Mornington | $6.559 \pm 0.029$ | $4.187 \pm 0.104$ | $1.995 \pm 0.357$ | $1.694 \pm 0.128$ |

**Table S5.** ANOVA results of soil properties and weather conditions among wine growing regions in 2017.

| | | $F_{(5,39)}$ | $p$ | Pr (>F) |
| --- | --- | --- | --- | --- |
| Soil | pH | 2.300 | 0.067 |  |
|  | C | 5.891 | 5.37E-04 | *** |
|  | N | 3.981 | 0.006 | ** |
|  | C:N | 6.309 | 3.27E-04 | *** |
|  | Nitrate | 11.640 | 1.56E-06 | *** |
|  | Ammonium | 1.780 | 0.144 |  |
| Weather | Mean solar radiation | 631.700 | <2e-16 | *** |
|  | Mean high temperature | 22.100 | 1.14E-09 | *** |
|  | Mean low temperature | 762.600 | <2e-16 | *** |
|  | Maximum temperature | 45.580 | 7.09E-14 | *** |
|  | Minimum temperature | 3823.000 | <2e-16 | *** |
|  | Mean temperature | 18.290 | 1.12E-08 | *** |
|  | Precipitation | 172.300 | <2e-16 | *** |
|  | Mean relative soil moisture | 0.901 | 0.492 |  |
|  | Mean evaporation | 0.729 | 0.607 |  |
|  | Mean transpiration | 188.100 | <2e-16 | *** |

**Figure S1.**
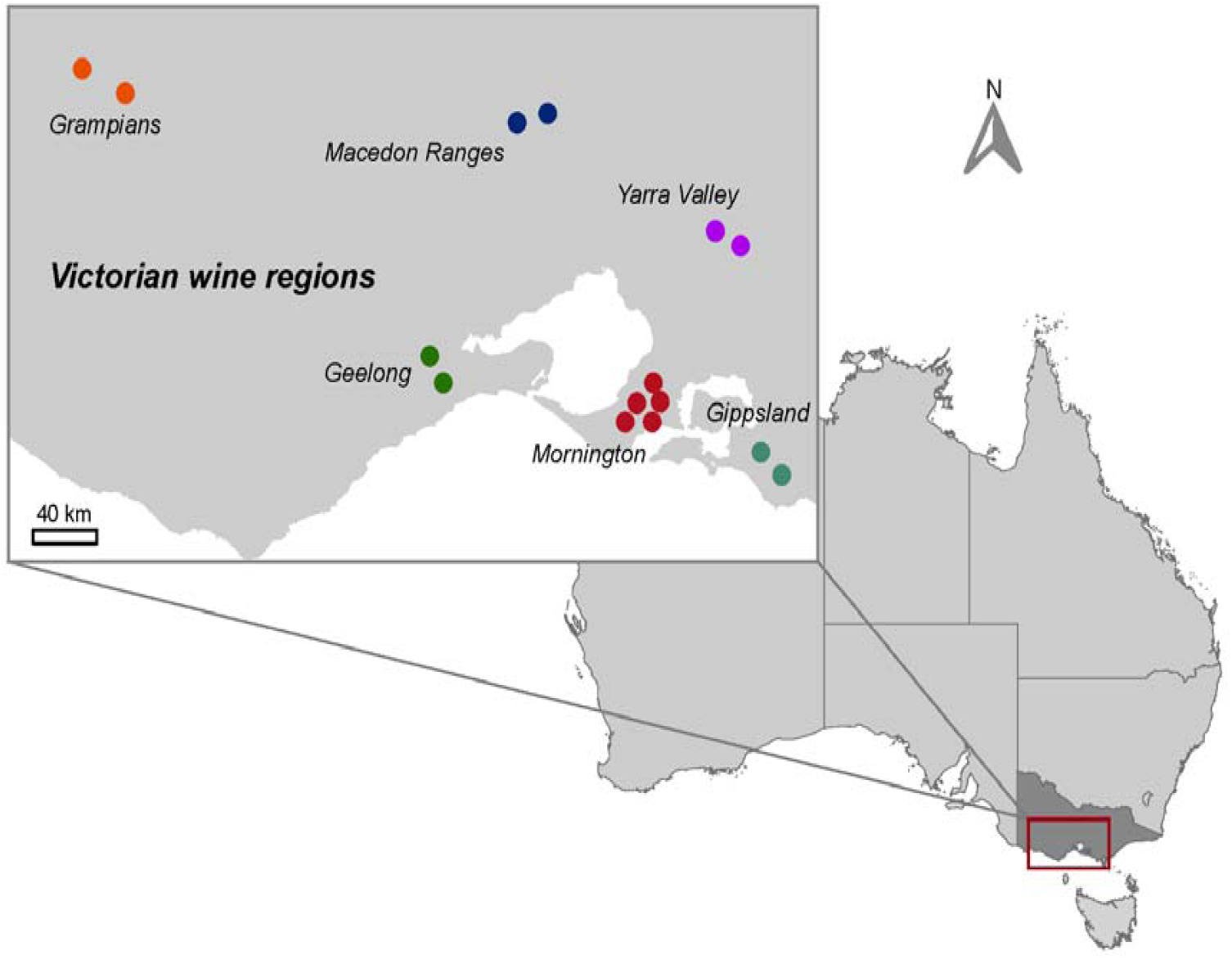
Map of 15 sampling vineyards from six Pinot Noir wine-producing regions in southern Australia, spanning 400 km (E-W).

**Figure S2.**
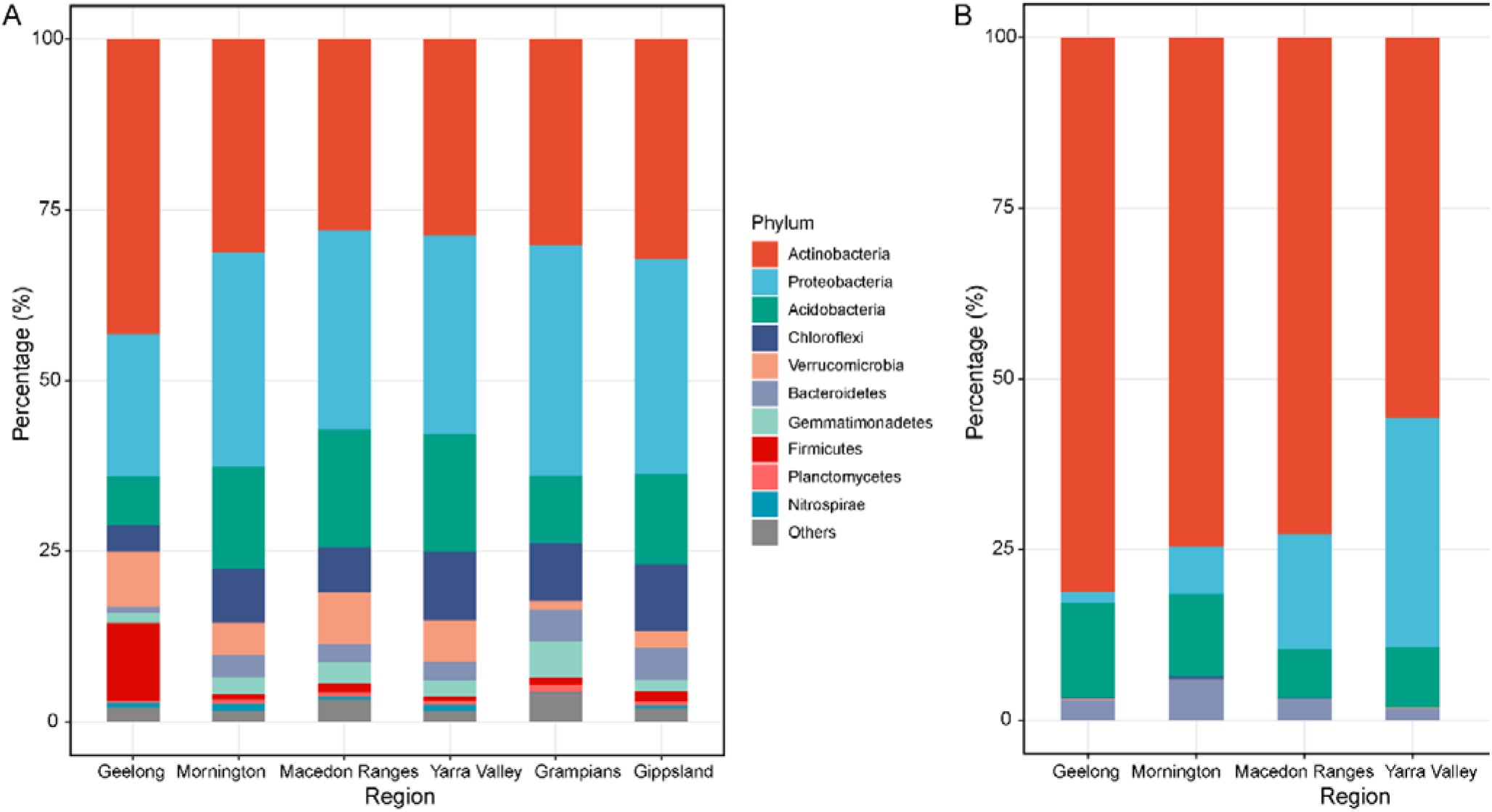
Vineyard soil microbial community compositions across all samples from six grape growing regions. Shown are average percentages of taxa (characterised to the phylum level) across sites in each region: (A) dominant bacterial phyla with greater than 1.0% relative abundance; (B) fungal phyla.

**Figure S3.**
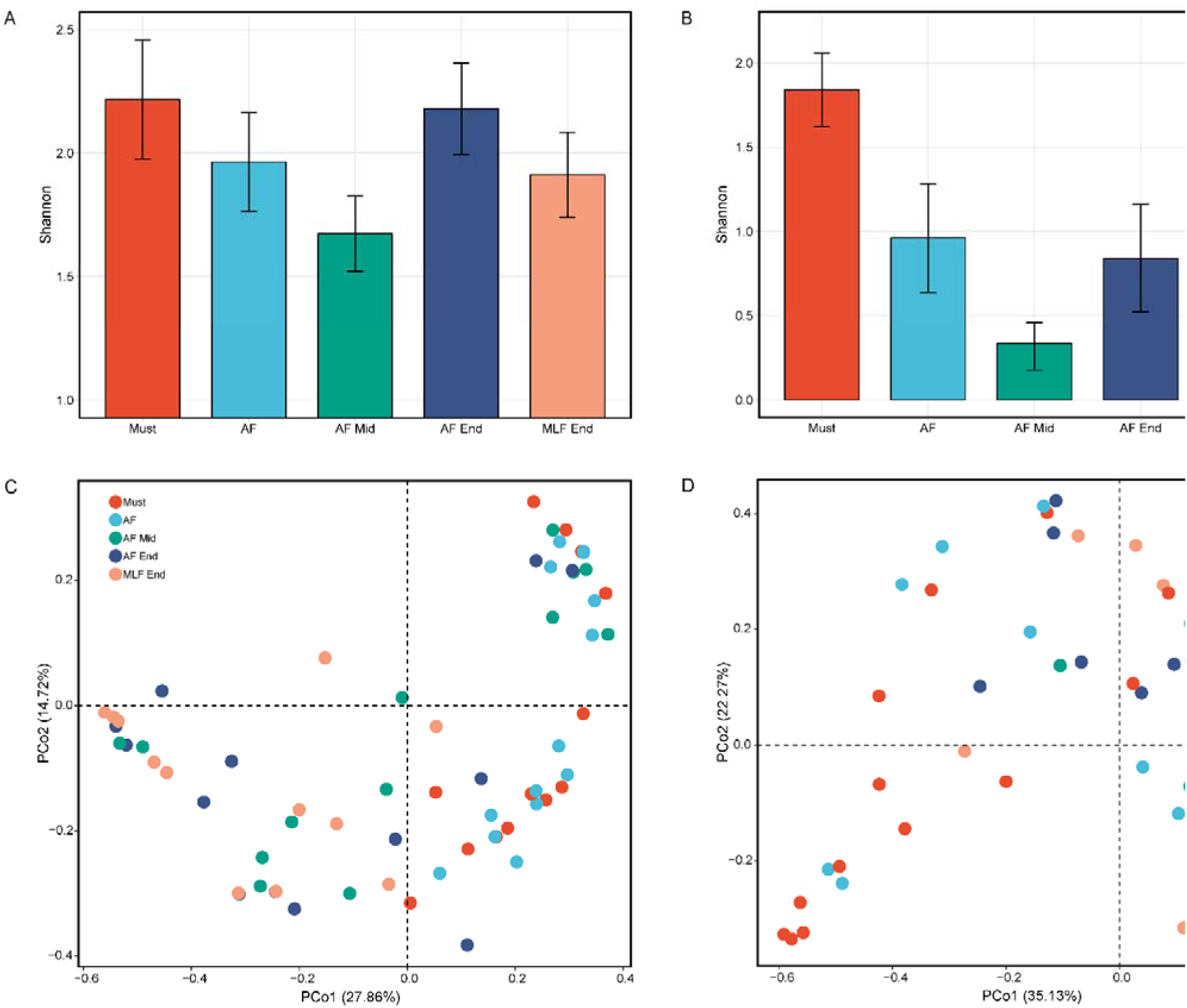
Stage of fermentation influences microbial diversities. Bacterial (A) and fungal (B) α-diversity shannon index) changes during wine fermentation. Bray-Curtis distance PCoA of bacterial communities (C) and fungal communities (D) according to the fermentation stage.

**Figure S4.**
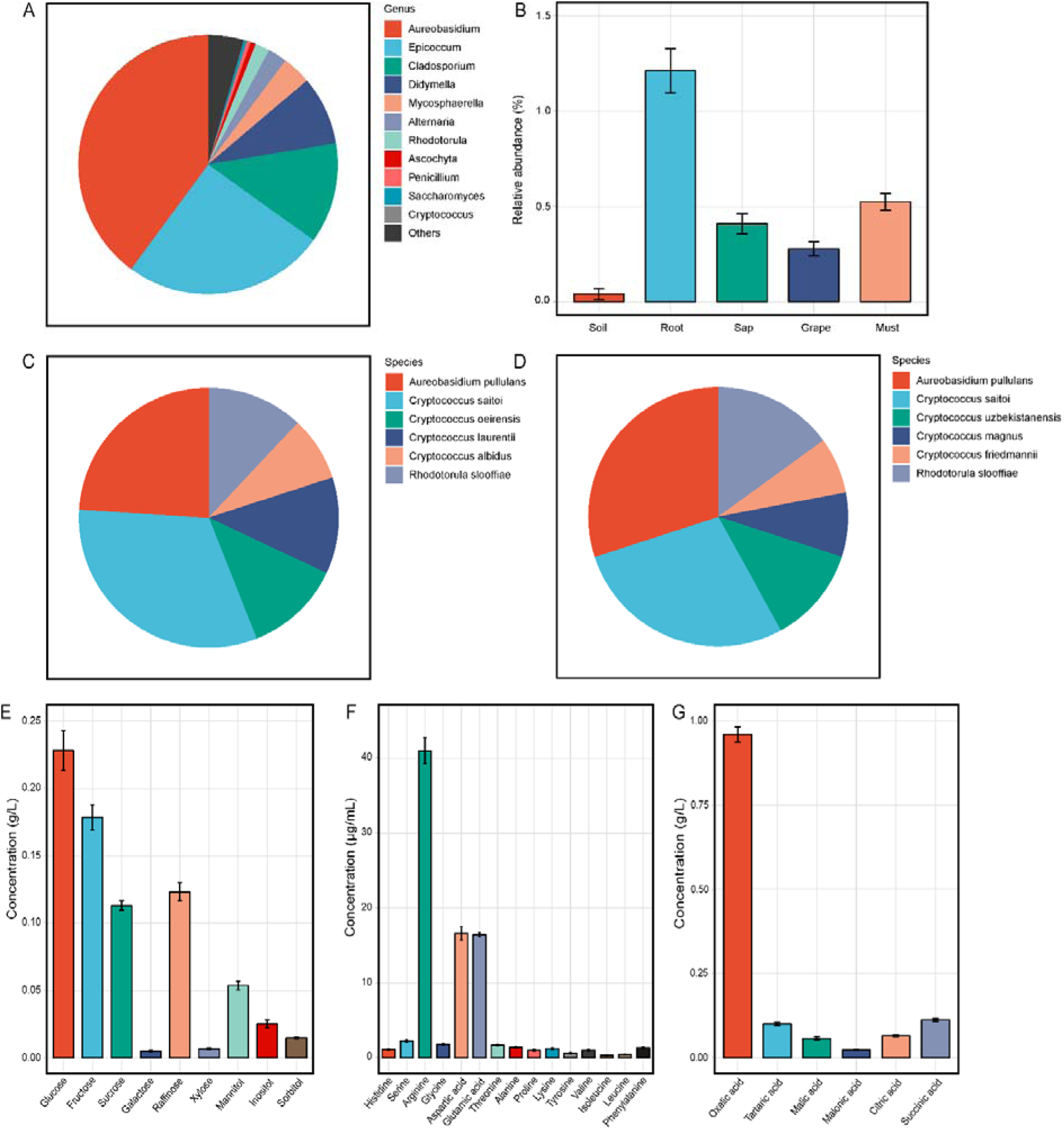
Xylem sap is a potential translocation medium of yeasts from the soil to the grapevine. (A) Fungal taxa in the genus level of xylem sap from the vineyard. (B) Relative abundances of *S. cerevisiae* in the soil, root, xylem sap, grape, must. Cultured yeast species from the xylem sap from vineyard (C) and glasshouse (D). Nutrient compositions of xylem sap: carbohydrates (E), total amino acids (protein and free) (F), and organic acids (G).

## Supplementary Materials and Methods

### Xylem sap collection

Vine shoots (n = 10) were harvested from the vineyard (Table S1) under aseptic conditions and then centrifuged (39). The shoots were peeled to remove the outer bark and phloem layer, sterilised with 70% ethanol and cut to fit into 10-ml sterile centrifuge tubes that had 10 sterile glass beads at the bottom. The shoots were centrifuged at 15,000 × *g* for 10 min at 4°C and the xylem sap collected giving ~ 2.0 mL per sample. Additional xylem sap (n = 5) were sampled from *Vitis vinifera* Shiraz grapevines grown in a glasshouse at the University of Melbourne. Xylem fluid was extracted with a pressure cylinder apparatus (similar to a Scholander pressure chamber) (83). Grapevines were uprooted and the soil was carefully removed from the roots. On an aseptic bench, the main trunk was cut 5-10 cm above the roots with a sterile blade, girdled to remove phloem tissue, sterilised by immersion in 70% ethanol, and immediately inserted into a pressure cylinder. The cylinder applied 60-70 kPa pressure for two hours to extract the xylem sap (~ 4.0 mL). Xylem sap was divided into two subsamples, one used immediately to isolate living yeasts and the other flash frozen with liquid nitrogen and stored at −80°C for DNA extraction, next-generation sequencing and chemical analysis.

### Chemical analysis of xylem sap composition

Carbohydrates of xylem sap were determined using enzymatic methods by Megazyme assay kits (Megazyme, Ireland) following the manufacturer’s protocol. Amino acids (free and protein-bound) were determined using pre-column derivatisation with 6-aminoquinolyl-N-hydroxysuccinimidyl carbamate followed by separation and quantification with the ACQUITY Ultra Performance LC (UPLC; Waters, MA, USA) system at the Australian Proteome Analysis Facility. The column was an ACQUITY UPLC BEH C18 column (1.7 μm × 2.1 mm × 5 mm) with detection at 260 nm (UV) and a flow rate of 0.7 mL/min at 57 – 60°C. Identification and quantitation of the amino acids was performed against a set of prepared standards, with DL-norvaline as the internal standard (84, 85). Organic acids were determined using a Waters High-performance liquid chromatography (HPLC) (Waters, MA, USA) based on Andersen et al (1989) (86) with modification. Here, 20 μL xylem sap was injected through a Synergi™ Hydro-RP LC Column (250 mm × 4.6 mm × 4 μm; Phenomenex Inc, CA, USA) at 60°C, with detection at 210 nm (UV). Mobile phases at a flow rate of 1.0 mL/min, with 20 mM potassium phosphate buffer (A, pH = 1.5) and 100% methanol (B) following a gradient programme: (0-2.5) min, 100% A; (2.5-2.9) min, linear ramp to 30% B; (2.9-8.0) min, 30% B; (8.0-8.5) min, linear ramp to 100% A; (8.5-10) min, 100% A. Identification and quantitation of the compounds was performed using a set of prepared standards.

### Isolation and identification of yeasts from xylem sap

Yeasts were isolated and identified from xylem sap to explore the potential translocation mechanism of yeasts in the vineyard. Xylem sap was serially diluted and plated (0.1 mL) onto solid yeast extract peptone dextrose (YPD) medium that was supplemented with 34 mg/mL chloramphenicol and 25 mg/mL ampicillin to inhibit bacterial growth. Plates were incubated at 28°C for 2-3 days in aerobic conditions. Single colonies with different morphological types were streaked onto Wallerstein Nutrient (WLN) agar media to obtain pure cultures. DNA was recovered from pure colonies using the MasterPure™ Yeast DNA Purification Kit (Epicentre, Madison, WI) following the manufacturer’s instructions. The 26S rDNA D1/D2 domain was amplified using primers NL1/4 (87) for sequencing by Australian Genome Research Facility (AGRF). Sequences were trimmed, aligned and analysed using BLAST at NCBI (http://blast.ncbi.nlm.nih.gov/Blast.cgi). Sequence data was uploaded to Genbank with accession numbers MN847682 - MN847692.

## Notes

#### Summary of Updates

Revisions for statistical analysis, proofreading, supplementary data added

